# Hip and knee kinematics display complex and time-varying sagittal kinematics during repetitive stepping: Implications for design of a functional fatigue model of the knee extensors and flexors

**DOI:** 10.1101/005538

**Authors:** Corey J. Scholes, Michael D. McDonald, Anthony W. Parker

**Affiliations:** Sydney Orthopaedic Research Institute, Chatswood, NSW; Institute of Health & Biomedical Innovation, Queensland University of Technology, Brisbane, Queensland, Australia; School of Exercise and Nutrition Sciences, Queensland University of Technology, Brisbane, Queensland, Australia

**Keywords:** variability, biomechanics, functional, velocity, displacement

## Abstract

The validity of fatigue protocols involving multi-joint movements, such as stepping, has yet to be clearly defined. Although surface electromyography can monitor the fatigue state of individual muscles, the effects of joint angle and velocity variation on signal parameters are well established. Therefore, the aims of this study were to i) describe sagittal hip and knee kinematics during repetitive stepping ii) identify periods of high inter-trial variability and iii) determine within-test reliability of hip and knee kinematic profiles. A group of healthy men (N = 15) ascended and descended from a knee-high platform wearing a weighted vest (10%BW) for 50 consecutive trials. The hip and knee underwent rapid flexion and extension during step ascent and descent. Variability of hip and knee velocity peaked between 20-40% of the ascent phase and 80-100% of the descent. Significant (p<0.05) reductions in joint range of motion and peak velocity during step ascent were observed, while peak flexion velocity increased during descent. Healthy individuals use complex hip and knee motion to negotiate a knee-high step with kinematic patterns varying across multiple repetitions. These findings have important implications for future studies intending to use repetitive stepping as a fatigue model for the knee extensors and flexors.

## Introduction

The progression and effect of muscle fatigue on knee function during locomotion remains poorly understood due to limitations in monitoring dynamic muscle fatigue. The experimental design and the model used to define and monitor fatigue is a critical factor in determining fatigue-related changes in muscle function. The model comprises the muscles of interest, the exercise protocol, the measures used to quantify fatigue, the timing of measurement and the operational definition of fatigue ^1^. The need to maximize external validity has prompted increased use of tasks that mimic occupational or sporting activities, such as jumping, squats or hopping ^2–4^. Stepping onto a step is a functional movement performed frequently in occupational situations ^5^. It requires high amounts of work performed by the hip and knee extensors to raise the body onto the raised surface. However, there is a lack of information regarding the internal validity of fatigue models involving functional tasks, such as stepping. Specifically, how effectively a task such as stepping induces fatigue in the quadriceps and hamstrings.

Monitoring fatigue onset and progression within the knee extensors and flexors during stepping is a complex undertaking. Surface electromyography (sEMG) has been used extensively for monitoring muscle fatigue in-vivo by detecting changes in the muscle activation signal ^6, 7^. In particular, the spectral shift of the sEMG signal to lower frequencies during static contractions is a valid and reliable measure of localised muscle fatigue ^8, 9^. However, neuromuscular changes in the sEMG signal are confounded by factors during dynamic movement such as variations in muscle length, muscle force output and contraction velocity ^8, 10, 11^. Importantly, joint kinematics are directly related to relative displacement of the surface electrodes and the underlying muscle fibres, which affects the signal properties in both time and frequency domains ^10, 12^. Previous studies have demonstrated a relationship between the variability of joint kinematics and the variability of the sEMG signal during dynamic movements ^12^. Therefore, sEMG analysis of muscle activity during dynamic, functional movement should be preceded by kinematic investigation to identify potentially variable periods.

A strategy to reduce the variability in the sEMG signal during dynamic movement is to select the most mechanically reproducible portion of a repetitive movement ^12, 13^. If it is assumed that confounding mechanical variables remain invariant from trial to trial, then the spectral changes observed in the sEMG signal can be related to physiological processes ^12^. While this strategy has been successfully demonstrated during lifting ^13–15^, there is little information on the variability of hip and knee kinematics during a stepping task. Hip and knee kinematics reflect knee extensor and flexor muscle length and contraction velocity, therefore a detailed kinematic analysis should precede any attempt to validate stepping as a functional fatigue model to identify periods of the movement that may be highly variable and thus unsuitable for sEMG analysis. The aims of this study therefore, were threefold: Firstly, describe the kinematics of the knee and hip during a repetitive stepping task. Secondly, identify the period of peak inter-trial variability of hip and knee kinematics during step ascent and descent. Thirdly, assess the within-session reliability of hip and knee kinematics during repetitive stepping.

## Methods

### Subjects

A sample of convenience comprising 15 healthy males (age: 20.7±2.5 yrs; height: 1.78±0.05 m; weight: 72.6±9.0 kg) were recruited from the university population to participate in the study. Males aged between 18-25yrs were tested to minimize age and gender effects on movement kinematics. All subjects participated in recreational sport at least twice a week. They were asked to avoid intense exercise the day prior to each test session and on test days. None of the participants were familiar with the stepping exercise prior to the experiment. Each volunteer indicated that they were unaffected by any musculoskeletal or neurological conditions that may have impaired their ability to perform the experimental tasks. The QUT Human Research Ethics Committee granted ethical approval for this study and written informed consent was obtained prior to testing.

### Study Design

This study was a part of a larger experiment to compare the efficacy and reliability of repetitive stepping and isokinetic exercise to induce muscle fatigue in the knee extensors. The experiment comprised of a test-retest cross-over study design, with each participant performing each fatigue protocol twice on separate days for a total of four test sessions (2 protocols X 2 tests). The sessions were conducted at the same time of the day in randomised order, with a minimum separation of 14 days between each test.

### Data Collection

The stepping protocol was performed in the motion analysis laboratory of the Institute of Health and Biomedical Innovation, QUT. Prior to the start of the test, reflective markers (10 mm) were placed in a modified lower-limb Helen Hayes marker set ^16^. Markers were placed bilaterally on the anterior superior iliac spines, the lateral femoral epicondyle, lateral malleolus of the fibula, calcaneus, and head of the second metatarsal, as well as the mid-sacrum. Markers mounted on wands were secured with Velcro straps to mid-thighs and mid-calf bilaterally. Three-dimensional coordinates of the markers were collected with a 6-camera system at 200Hz (VICON Motion Systems Ltd., Oxford, UK). Kinetic data was collected at 1000Hz with a force-plate (OR6-2000, AMTI, USA) mounted on a frame bolted to the floor of the laboratory (Figure 1). The contralateral foot made contact with forceplate mounted in the floor of the laboratory and a pressure-switch on the frame-mounted plate to identify the ascent and descent phases of the stepping movement while the lead foot remained stationary on the frame-mounted plate. Analog and video data were captured synchronously with motion capture software (VICON Motus 9.2, VICON Motion Systems Ltd., Oxford, UK) and stored on a computer.

**Figure 1.**
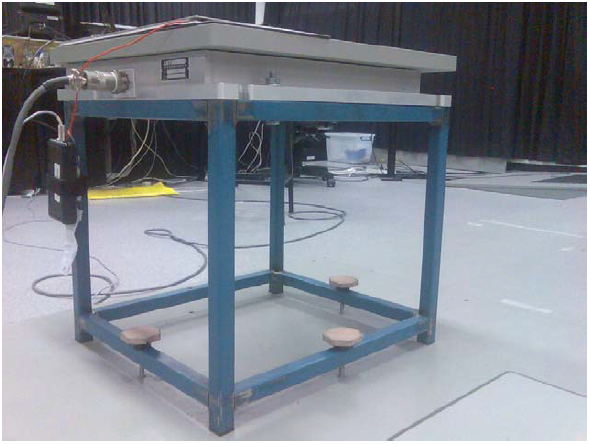
Forceplate fixed to a frame to provide a stepping platform for the repetitive stepping task. Note the pressure switch on the top of the platform with battery pack attached with red wire (left side of frame)

The stepping protocol involved 50 stepping trials performed as rapidly as possible, while the participant wore a vest containing additional load equal to 10% of their bodyweight. A step trial involved the participant raising themselves to an upright standing posture on the frame (ascent) before returning their trailing limb to the floor-mounted force plate (descent). At the start of the test, the subject placed the leading test leg on the frame mounted platform matched to the height of the lateral tibial condyle (45 – 55cm high), with the contralateral leg placed shoulder width apart on the floor platform. All participants were tested with the right leg leading. The start and end postures were demonstrated to the subject prior to the commencement of each session to standardize the movements. Participants performed 10 trials at a comfortable pace to familiarise themselves with the task. Participants were encouraged to attain a straight knee of the trail limb before the start of each stepping trial and a straight posture of both knees after ascent onto the frame (Figure 2). In addition, the lead foot was placed in a central point on the frame mounted platform in the anterior-posterior axis, offset to the right of the midline. The participant was asked to maintain this position of the leading leg, including contact with the platform throughout the test. Following the familiarisation period, the participant performed the series of 50 step trials as fast as possible. Arm position was not constrained during the experiment.

**Figure 2.**
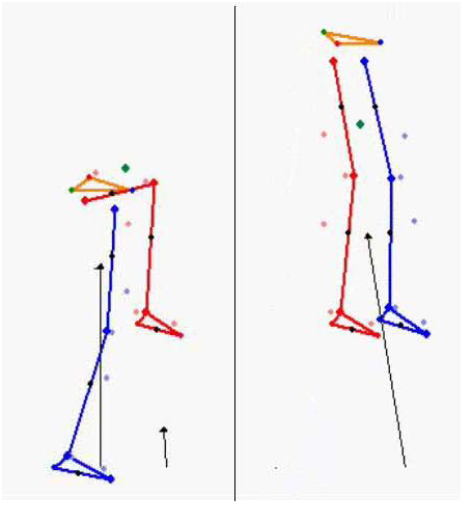
Key postures of the stepping trial illustrated with ground reaction force vectors. The end position of the ascent movement (right) was also the starting position for the step-down movement

### Data Analysis

A custom-written function in Matlab (version 2007a, Mathworks Inc. USA) split the stepping movement into ascent and descent phases based on the data collected from the pressure switch and the floor-mounted forceplate. The beginning of the ascent phase was defined as the point at which the trail limb left the floor-mounted forceplate minus an offset. The offset was calculated as the average time between the trail foot impact at the end of the descent and the foot leaving the plate to ascend the platform. The end of the ascent phase was defined as the point at which the trail limb contacted the pressure-switch mounted on the forceplate mounted on the platform. The descent phase was defined as the period between the foot leaving the pressure-switch and contacting the floor forceplate.

Hip and knee flexion displacement and velocity were calculated during ascent and descent phases using established equations ^16^. The data were first resampled to 1000Hz using a quintic spline processor for synchronization with the analog data. Following phase identification, each data vector was then interpolated to a length of 100 points using a fast Fourier transform (FFT) interpolation function ^17^. Key variables extracted from each phase (ascent and decent) were maximum and minimum flexion angles, range of motion and timing of maximum joint flexion. The magnitude and timing of peak velocity in flexion and extension were determined. All timings were expressed as % of the phase duration.

### Statistical Analysis

Data was assessed for normality and equality of variance prior to further analysis. Data from the 50 stepping trials were grouped evenly into 10 blocks of 5 trials, with inter-trial variability of angular displacement and velocity calculated over each 5-trial block with root mean square error (RMSE). Inter-joint differences in kinematics were assessed at trial block 1 and trial block 10 with Mann-Whitney U tests. Ten RMSE vectors were calculated for each variable for each participant, for the ascent and descent phases of stepping. Calculating the error relative to the mean of each trial block accounted for any changes in kinematic parameters across the 50-trial test. Kruskal-wallis ANOVA with Dunn-Sidak post-hoc comparisons to assess within-session changes in angular displacement and velocity across trial blocks (Blocks 1-10). Significance level was set a-priori at P < 0.05 for all statistical tests, which were performed in Minitab (version 16, Minitab Inc, MA, USA). Linear changes in kinematic variables between trial blocks were quantified with linear regression in Microsoft Excel (v2010).

### Results

All subjects successfully completed both sessions of 50 stepping trials. Angular displacements of the hip and knee were similar and both demonstrated a pattern of flexion-extension during ascent and the reverse during descent (Figure 3). During the first block of trials, the hip and knee moved through similar (P = 0.097) ranges of motion and finished the movement with similar minimum flexion angles during step ascent (Table 1). In contrast, peak flexion velocity of the knee during ascent was significantly (P < 0.01) higher than the hip, while no significant difference was observed for peak extension velocity (Table 1). The peak flexion velocity of the knee during descent was significantly greater (P < 0.01) and occurred later in the movement than the hip (Table 1). Inter-joint differences in kinematics remained during the last blocks of trials (Table 2). However, the peak extension velocity of the knee during ascent was significantly greater than the hip during the last trial block, although no difference was detected during the first trial block.

**Figure 3.**
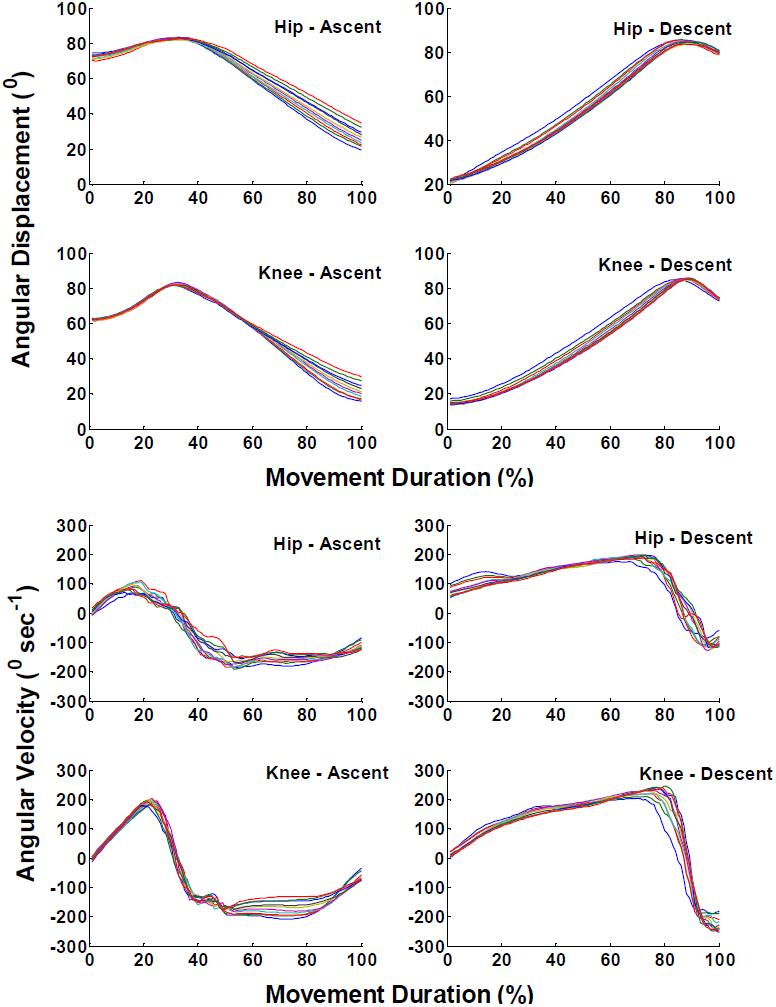
Average angular displacement (top) and angular velocity (bottom) curves across trial blocks (10) for the hip and knee during the ascent and descent phases of the repetitive stepping movement

**Table 1.** Hip and knee kinematics (median and inter-quartile ranges) with inter-joint differences during Trial Block 1 of the repetitive stepping task

|  | Hip | Knee | P-value |
| --- | --- | --- | --- |
| **Ascent** |  |  |  |
| Range of motion | 67.7 (63.3 – 78.3) | 73.4 (68.5 – 77.7) | 0.097 |
| Minimum Joint Angle | 22.1 (11.4 – 26.5) | 16.2 (11.6 – 20.9) | 0.097 |
| Peak Flexion Velocity | 164.2 (142.6 – 205.5) | 251.2 (227.9 – 276.6) | <0.01 |
| Peak extension velocity | 225.2 (206.9 – 272.6) | 254.8 (237 – 292.3) | 0.051 |
| **Descent** |  |  |  |
| Peak Flexion Velocity | 203.4 (190.2 – 212.7) | 246.2 (226.7 – 276.9) | <0.01 |
| Peak Flexion Velocity Timing (%) | 70 (62-83) | 84 (74-87) | 0.049 |

**Table 2.** Hip and knee kinematics (median and inter-quartile ranges) with inter-joint differences during Trial Block 10 of the repetitive stepping task

|  | Hip | Knee | P-value |
| --- | --- | --- | --- |
| **Ascent** |  |  |  |
| Range of motion | 52.3 (46.5 – 75.7) | 60 (53.3 – 67.8) | 0.41 |
| Minimum Joint Angle | 37.4 (14.3 – 43.4) | 29.6 (22.0 – 36.6) | 0.36 |
| Peak Flexion Velocity | 175.2 (153.1 – 187.6) | 288 (277 – 315.3) | <0.01 |
| Peak extension velocity | 193.4 (167 – 234.7) | 276.7 (259.1 – 310.7) | <0.01 |
| **Descent** |  |  |  |
| Peak Flexion Velocity | 209.8 (183.5 – 254.7) | 304.7 (250.4 – 322.7) | <0.01 |
| Peak Flexion Velocity Timing | 82 (72 – 87) | 86 (85 – 90) | 0.018 |

### Discussion

The first aim of this study was to describe the kinematics of the knee and hip during a repetitive stepping task. On average, healthy young men employed a pattern of hip-knee flexion followed by rapid extension to ascend the knee-high step and a reverse pattern to descend the step. While this is the first study to report hip and knee kinematics with such a high step rise (∼50cm), the angular displacements observed were comparable to previous studies at lower step heights (18cm) ^18, 19^. The flexion-extension joint motion during step ascent may reflect the participants’ attempts to take advantage of the stretch-shorten cycle to generate adequate joint power ^20^. That is, initial stretching of the hip and knee extensors during joint flexion enhance the mechanical output of the proceeding concentric muscle contraction as the hip and knee extend during step ascent. However, inter-joint differences in peak joint velocity and peak joint velocity timing revealed a more complex movement pattern than initially thought. These complex patterns may be explained by the energy requirements of raising and lowering the body’s centre of mass during the stepping motion ^21^, which may require redistribution of forces between the hip and knee extensors ^22^. The kinematics observed are more complicated than the pure joint extension during ascent and flexion during descent that would be assumed to occur. Future studies should consider partitioning the ascent and descent phases of the stepping movement into sub-phases representing eccentric and concentric muscle actions.

The second aim of the study was to identify the period of peak inter-trial variability of hip and knee kinematics during step ascent and descent. Inter-trial variation in joint kinematics has important implications for sEMG of the hip and knee extensors during stepping. In particular, sEMG signal analysis can be confounded by changes in movement biomechanics (non-physiological factors), which interfere with the observation of physiological processes, such as muscle fatigue (De Luca, 1997). Strategies to minimise confounding by variations in joint kinematics have been proposed which involve selecting the most mechanically reproducible part of a cyclic movement ^12^. The presence of a stretch-shorten pattern during both phases of the stepping trial produced spikes in inter-trial variation of joint velocity coinciding with the transition from flexion to extension. Hip and knee angular velocity RMSE nearly doubled during the transition periods compared to the rest of the movement phase. This can be attributed to variations in the timing of the transition between trials. That is, differences in timing of as little as 1% between trials would mean that positive (flexion) velocities were averaged with negative (extension) joint velocities. Therefore, future studies that conduct sEMG of functional protocols should proceed with caution when attempting to analyse signals collected from 21-40% of the step ascent and 81-100% of the descent, where flexion-extension transitions occur.

The third aim of the present study was to assess the within-session reliability of hip and knee kinematics during repetitive stepping. A key finding was that angular displacement and peak joint velocity did not remain stable across the 50-trial test. Such a result is not unexpected considering the fatiguing nature of the task, with reductions in joint range of motion, peak hip extension velocity and increased peak knee extension velocity possibly reflecting adaptation strategies to maintain maximum movement speed across the 50 trials ^23, 24^. However, these alterations may confound future sEMG analyses if stepping is to be used a functional fatigue model of the hip and knee extensors. In particular, estimates of sEMG signal frequency may be influenced by the changes in joint angle during ascent, as well as alterations in joint velocity during ascent and descent ^10, 11^. Knee kinematics are influenced by external constraints ^25^, with improved within-session reliability of knee kinematics during constrained squats compared to free squats and wall slides ^26^. These findings have important implications for future studies intending to use repetitive stepping as a fatigue model for the knee extensors and flexors, which may rely on surface EMG. Therefore future work may be required to constrain hip and knee motion with mechanical devices rather than verbal encouragement to improve reliability, as well as develop algorithms to adjust the sEMG analysis to compensate for changes in joint range of motion and velocity.

As with any study of this kind, the results should be interpreted in the context of its limitations. Firstly, the analysis was impeded by considerable between-participant variation of hip and knee kinematics, which is reflected in the inter-quartile ranges of the results (Figures 4–7). Study recruitment was restricted to a narrow age range of university males to minimize gender and age effects and attempts were made to standardize the stepping task for each participant, such as matching the step height to the tibial tuberosity, standardizing placement of the lead foot on the step, implementing familiarization trials and verbally encouraging the participant to attain standard postures. Despite these measures, the sample displayed considerable between-participant variability in hip and knee kinematics, which is a recognized characteristic of human movement ^27^, but also encourages the need for caution when interpreting the group results. The second limitation of the present study was the restriction of the analysis to the sagittal plane, with potential implications for sEMG analysis with respect to the secondary axes of motion at both joints. Previous work has illustrated that constraining lower limb posture with mechanical means can also improve tibial rotation reliability during squats ^26^. These findings provide opportunities to plan and execute dynamic sEMG analysis of functional, cyclical lower-limb movements to study muscle function.

**Figure 4.**
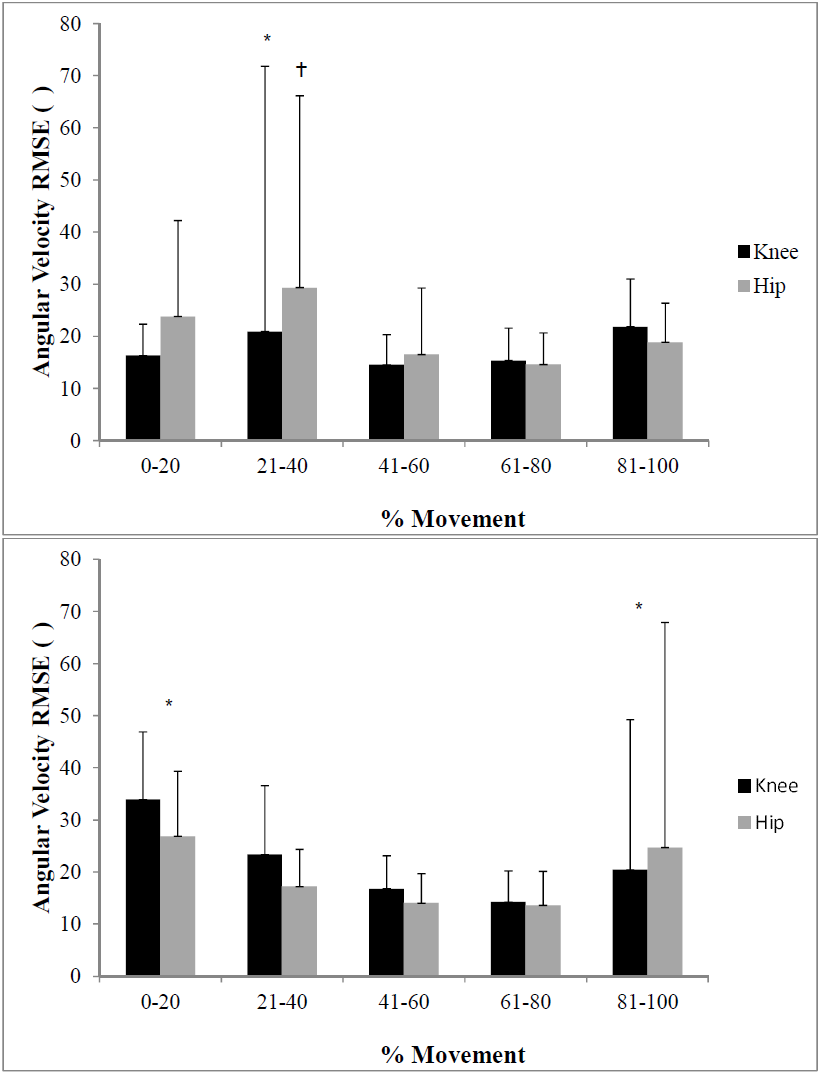
Inter-trial variability of angular velocity for the hip and knee compared across sub-phases during the ascent (top) and descent (bottom) of the stepping movement. * significantly (p < 0.05) different to all other sub-phases. † significantly different to 41-100%

**Figure 5.**
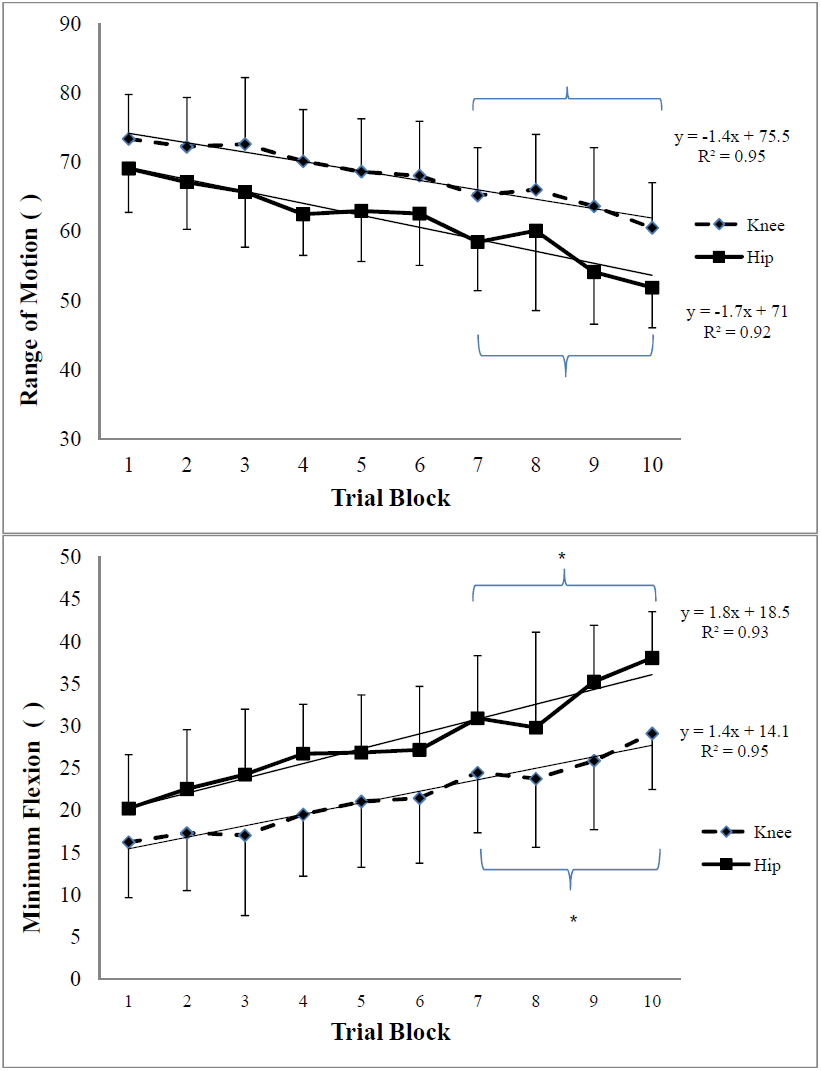
Within-test variation of angular displacement during step-up. Changes in range of motion (top) and minimum flexion (bottom) across trial blocks. Data represented by median ± inter-quartile range. *significantly different (p < 0.05) to blocks 1-3

**Figure 6.**
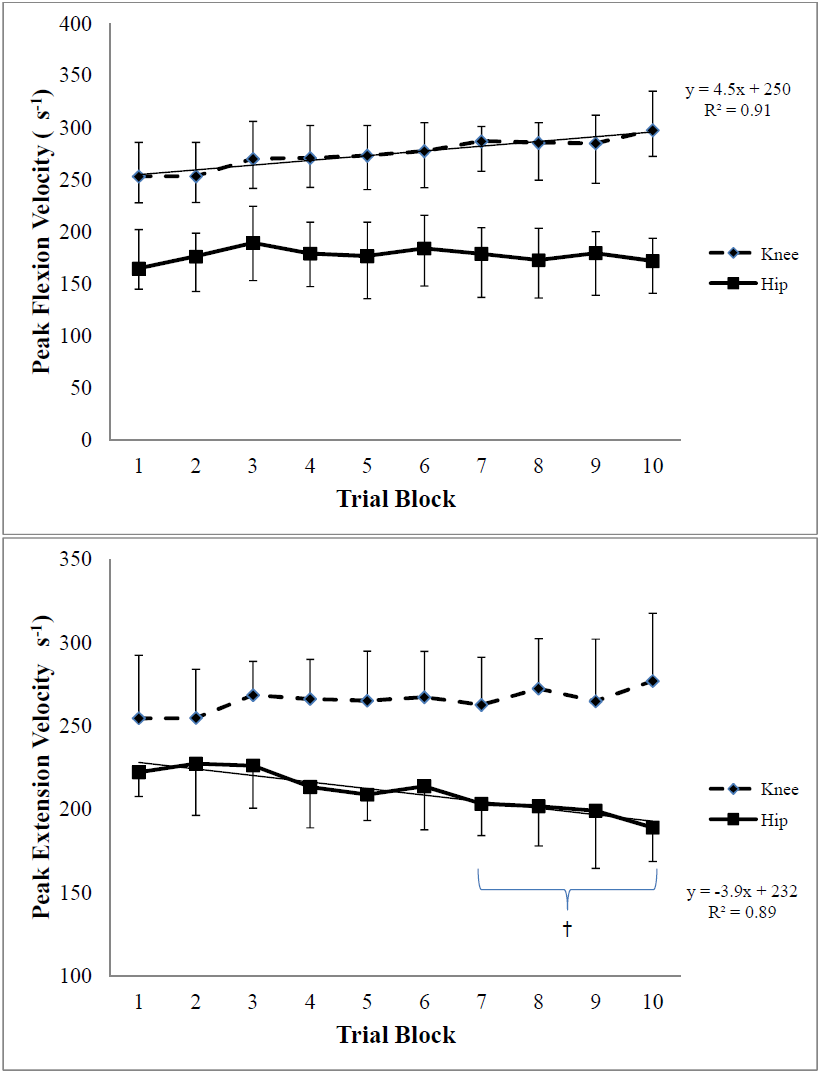
Within-test variation during step ascent of peak flexion velocity (top) and peak extension velocity (bottom) across trial blocks. Data represented by median ± IQR. * significantly different (p < 0.05) to trial blocks 1 and 2 † significantly different to trial blocks 1-3

**Figure 7.**
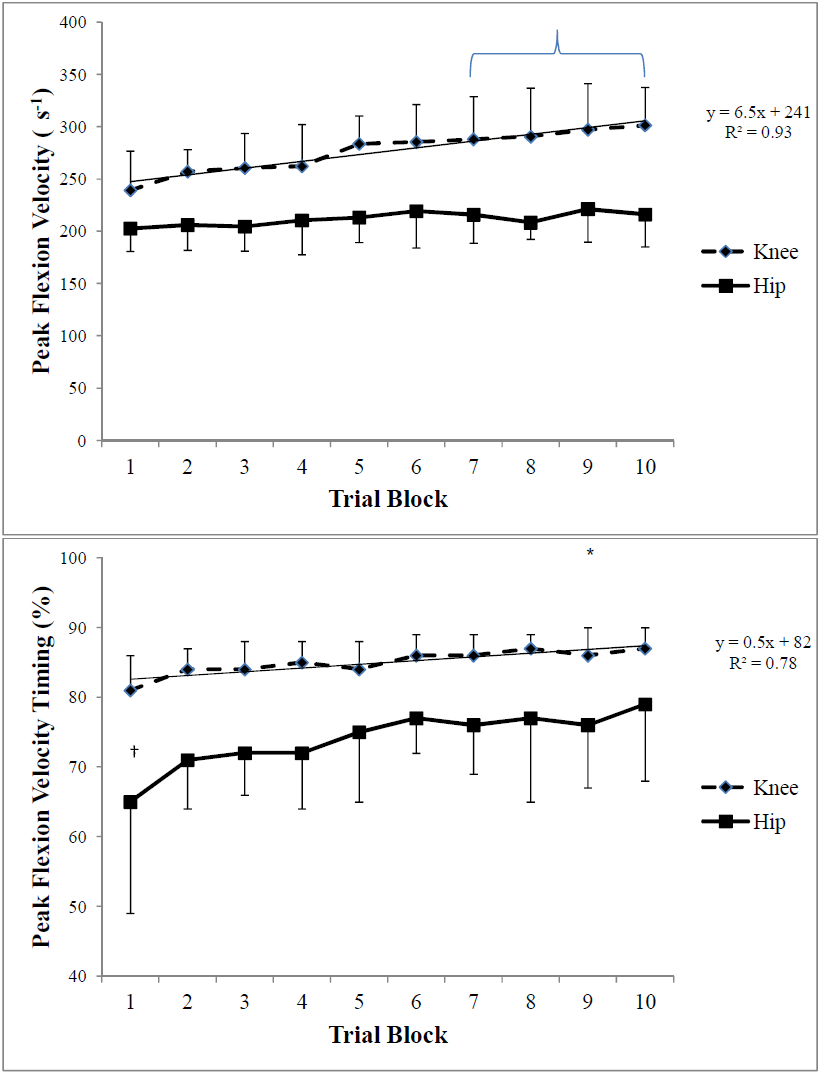
Within-test variation during step descent of peak flexion velocity (top) and time of peak flexion velocity (bottom) across trial blocks. Data represented by median±IQR *significantly different (p < 0.05) to blocks 1-3. † significantly different (p < 0.05) to all other blocks

### Conclusions

The hip and knee undergo complex patterns of sagittal motion to ascend and descend a knee-high step. Transitions between flexion-extension for the knee and hip during step ascent and descent coincide with periods of high inter-trial variability of joint kinematics which should be avoided in future sEMG analyses. Furthermore, hip and knee kinematics do not remain stable during a repetitive stepping task, possibly due to compensation strategies, which may further confound attempts to monitor muscle function. Future work should endeavor to constrain lower limb knee motion within the context of a functional lower-limb task to improve the reliability of sEMG data, as well as develop algorithms to adjust the signal to account for biomechanical variations within and between participants.

## Acknowledgements

The authors wish to acknowledge the technical assistance of Mr Alan Barlow and Dr Nathan Stevenson in the laboratory setup and data analysis of this work. Financial support for this study was provided by the Institute of Health and Biomedical Innovation, Queensland University of Technology. Dr Scholes was supported by the Sydney Orthopaedic Research Institute during the writing of this paper.

